# A-Prot: Protein structure modeling using MSA transformer

**DOI:** 10.1101/2021.09.10.459866

**Authors:** Yiyu Hong, Juyong Lee, Junsu Ko

## Abstract

In this study, we propose a new protein 3D structure modeling method, A-Prot, using MSA Transformer, one of the state-of-the-art protein language models. For a given MSA, an MSA feature tensor and row attention maps are extracted and converted into 2D residue-residue distance and dihedral angle predictions. We demonstrated that A-Prot predicts long-range contacts better than the existing methods. Additionally, we modeled the 3D structures of the free modeling and hard template-based modeling targets of CASP14. The assessment shows that the A-Prot models are more accurate than most top server groups of CASP14. These results imply that A-Prot captures evolutionary and structural information of proteins accurately with relatively low computational cost. Thus, A-Prot can provide a clue for the development of other protein property prediction methods.

## Introduction

Modeling the 3D structure of a protein from its sequence has been one of the most critical problems in biophysics and biochemistry (Kwon et al., 2021; Pereira et al., 2021). The knowledge of the 3D structure of a protein facilitates the discovery of novel ligands, function annotation, and protein engineering. Due to its importance, the community of computational protein scientists have been developing various prediction methods and assessed their performance in the large-scale blind tests, CASPs, which have continued for over two decades (Pereira et al., 2021). In CASP14, Deepmind demonstrated that their deep learning-based model, AlphaFold2 (AF2), predicts the 3D structures of proteins from their sequences with extremely high accuracy, comparable to experimental accuracy (Jumper et al., 2021). The source code of AF2 became publicly available recently, and many model structures of genes of biologically important organisms have been released (Jumper et al., 2021).

The success of AF2 can be attributed to the accurate extraction of coevolutionary information from multiple sequence alignments (MSA). The idea of using coevolutionary information for contact prediction or structure modeling has been widely used (de Juan et al., 2013; Hopf et al., 2014; Jones et al., 2015; Lunt et al., 2010; Marks et al., 2012a, 2012b; Ovchinnikov et al., 2014; Seemayer et al., 2014; Tillier & Charlebois, 2009; Weigt et al., 2009). However, previous attempts were not successful enough because coevolutionary signals in MSAs were not strong enough or too noisy. Various statistical mechanics-based models were proposed, but their accuracy, discriminating true contacts from false contacts, was limited. In CASP13, AlphaFold and RaptorX used a neural network to extract coevolutionary information from MSAs (Senior et al., 2020; Xu, 2019; Xu & Wang, 2019). They achieved significant improvements than previous CASPs, but their predictions were not accurate enough to obtain model structures comparable to experiments. Finally, in CASP14, with the help of the attention algorithm, truly accurate extraction of actual residue-residue contact signals from MSAs becomes available (Jumper et al., 2021).

Despite this remarkable achievement of AF2, certain limitations remain for this technology to be widely accessible and used for further development. First, the source code for training AF2 is not open-sourced yet. In the first release of AF2, only the production, model generation, part of AF2, and the model parameters are open. Second, training the AF2 architecture is computationally expensive. It is reported that Deepmind used 128 TPUv3 cores for approximately one week and four more days to train AF2. This amount of computational resources are not readily accessible for most academic groups. Thus, developing lighter and general models is still necessary.

To extract evolutionary information from MSAs, various protein language models have been proposed (Bepler & Berger, 2021; Madani et al., 2020; Rao, Liu, et al., 2021; Rao, Meier, et al., 2021; Rives et al., 2021; Strodthoff et al., 2020; Vig et al., 2020). Rao et al. proposed the MSA transformer model, an unsupervised protein language model using the MSA of a query sequence instead of a single query sequence. The model uses row and column attention of input MSAs and masked language modeling objectives. It is demonstrated that the model successfully predicted long-range contacts between residues. In addition, the model predicted other properties of proteins, such as secondary structure prediction and mutational effects, with high accuracy (Rives et al., 2021). These results indicate that the MSA Transformer model extracts the characteristics of proteins from their MSA profiles efficiently.

This study developed a new protein 3D structure prediction method, A-Prot, using MSA Transformer (Rao, Liu, et al., 2021). For a given MSA, we extracted evolutionary information with MSA Transformer. The extracted row attention map and input features were converted to a 2D residue-residue distance map and dihedral angle predictions. We bench-marked the 3D protein structure modeling performance using the FM/TBM-hard targets of CASP13 and 14 (Kinch et al., 2019, 2021). The results show that A-Prot outperforms most top server groups of CASP13 and 14 in terms of long-range contact predictions and 3D protein structure modeling.

## Methods

### Overview

The overall pipeline of the proposed protein 3D structure prediction method is shown in Figure. 1. We mainly combine the works of the MSA Transformer (Rao, Liu, et al., 2021) and the trRosetta (Yang et al., 2020). Given an MSA, it will input to the MSA Transformer to output MSA features and row attention maps. After a series of transformations like dimension reduction and concatenation etc., these two kinds of features will be transformed and combined into 2D feature maps, which is suitable for input to the trRosetta to finally output protein 3D structure.

**Figure. 1.**
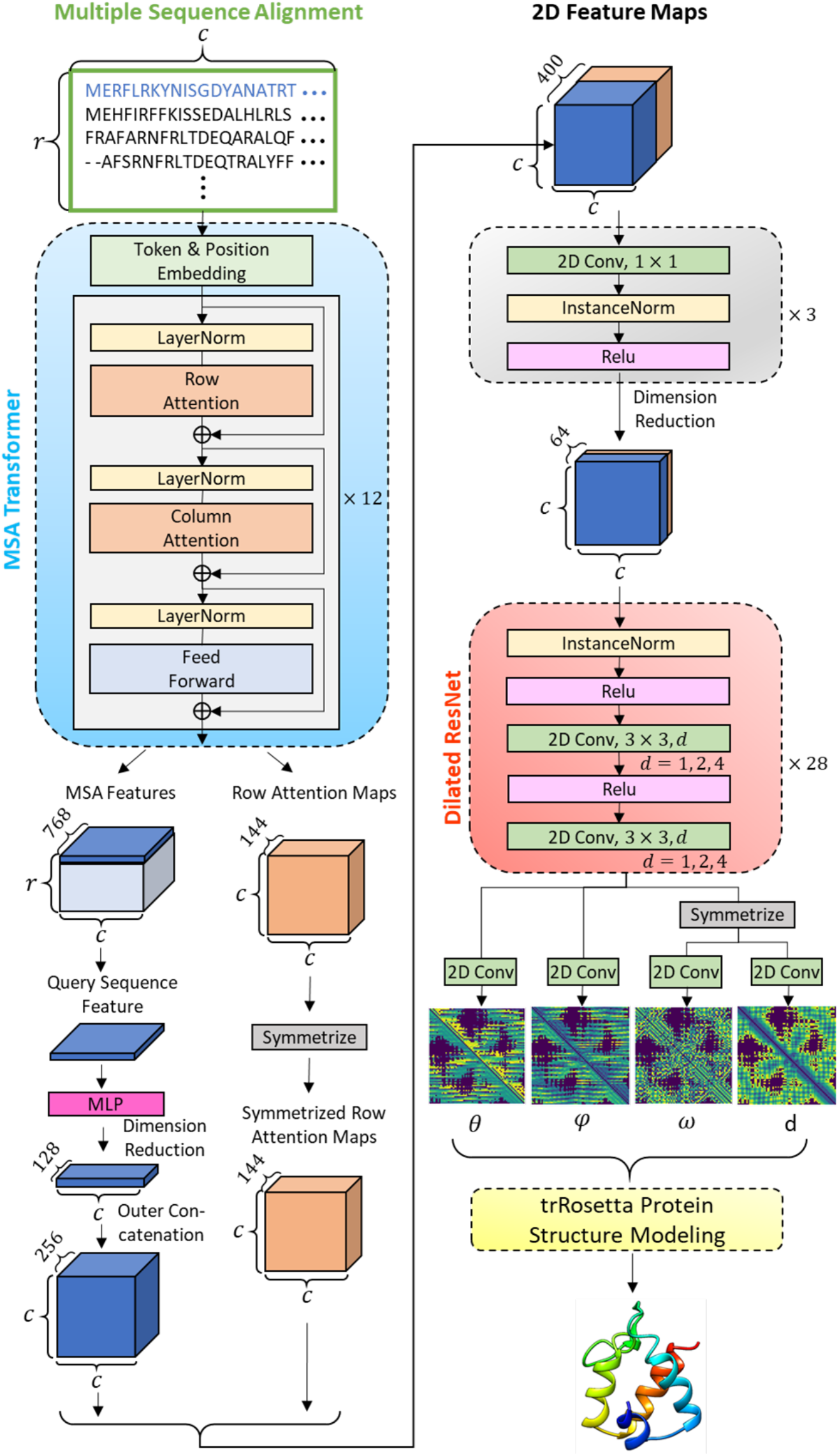
Pipeline of the proposed protein 3D structure prediction method. Input an MSA to the MSA Transformer to extract MSA Features and row attention maps. Then, the MSA features corresponding to the query sequence and the row attention maps are combined to a 2D feature maps by a set of transformations. Next, the 2D feature maps are input to a Dilated ResNet after dimension reduction to output inter-residue geometries, which further input to the trRosetta Protein Structure Modeling to output a predicted protein 3D structure.

The MSA Transformer used in this paper was an already pre-trained version that was learned from 26 million MSAs. It plays a role as a feature extractor that inputs an MSA and outputs its relative features.

trRosetta consists of a deep neural network part and protein structure modeling part. The deep neural network is modified to receive the MSA features from the MSA Transformer instead of the original manually engineered features generated by the statistic approach. Nevertheless, the protein structure modeling part remains the same.

### Dataset and MSA Generation

We used the procedure with MSA transformer to generate the MSAs of target sequences (Rao, Liu, et al., 2021). The MSA of a query sequence was generated using HHblits ver. 3.3.0 together with the unicluster30_2017_10 (Mirdita et al., 2017) and BFD (Steinegger et al., 2019) databases. If the number of detected homologous sequences exceeds 256, up to 256 sequences were selected through diversity minimization (Rao, Liu, et al., 2021).

To perform contact predictions, a customized non-redundant protein structure database and we call it PDB30. The PDB30 dataset consists of protein structures deposited in PDB before Apr-30-2018 whose resolution is higher than 2.5 A and sequence length is longer than 40 amino acids. Using 102,300 sequences that satisfy the condition, clustering analysis was performed using MMSeq2 (Steinegger & Söding, 2017) with a threshold of sequence identity of 30%, leading to 16,612 non-redundant sequences.

### Network Architecture

Let us define an input MSA as an *r* × *c* character matrix, where *r* and *c* are rows (number of sequences) and columns (sequence length) in the MSA, respectively. Through the token and position embedding of the MSA Transformer, the input MSA is embedded into a *r* × *c* × 768 tensor which is the input and output of each attention block. The attention block is composed in the order of row attention layer, column attention layer, and feed-forward layer. A layer normalization operation is followed by each layer. And each attention layer has 12 attention heads. The MSA Transformer is a stack of 12 such attention blocks.

Two kinds of features were extracted from the MSA Transformer to construct 2D feature maps by a series of transformations. (1) One is the last attention block’s output, a *r* × *c* × 768 tensor; we named it MSA features (Figure. 1). Only features corresponding to the query sequence are selected, which is a 1 × *c* × 768 tensor. Then, the dimension of this feature is reduced to 128 by an MLP (multi-layer perceptron) consist of 3 linear layers with neuron sizes 384, 192, 128. The dimension reduced 1D feature is then outer concatenated (redundantly expanding horizontally and vertically and then stacked together) to form a query sequence feature of a *c* × *c* × 256 tensor. (2) The other one is row attention maps that are derived from each attention head of each row attention layer, totally 12 × 12 = 144 attention maps stacked to shape in *c* × *c* × 144. Then it is symmetrized by adding it to its transposed tensor to yield symmetrized row attention maps. The query sequence feature and the symmetrized row attention maps are concatenated in the feature map dimension to form a 2D feature map that is a *c* × *c* × 400 tensor.

The dimension of the 2D feature maps afterward reduced to 64 by three convolutional layers that consist of 256, 128, 64 kernels of size 1 × 1 (Lin et al., 2013), where each convolutional layer is followed by an instance normalization and a ReLU activation. Then this *c* × *c* × 64 tensor is input to the Dilated ResNet consist of 28 residual blocks (He et al., 2016), each having one instance normalization, two ReLU activations, and two convolutional layers each with 64 kernels of size 3 × 3. Dilation is applied to the convolutional layers cycle through the residual blocks with rates of 1, 2, 4. After the last residual block, there are four independent convolutional layers with 25, 13, 25, 37 kernels of size 1 × 1, each for predicting inter-residue geometries of *θ, φ, ω*, d, respectively. The values for the inter-residue geometries are all discretized into bins; each bin is treated as a classification label. Please refer to the reference (Yang et al., 2020) for more details about the inter-residue geometries as we used the same settings. Finally, the trRosetta Protein Structure Modeling module will predict and modeling the protein 3D structure based on the inter-residue geometries information.

### Training and Inference

At the training stage, we fixed the parameters in the MSA Transformer. In contrast, parameters in the other deep neural networks were trained with a batch size of 16 with gradient accumulation steps, a learning rate of 1e-3, using the RAdam optimizer (Liu et al., 2020), the cross-entropy loss was calculated with equal weight for the four inter-residue geometry objectives. The total model was trained end-to-end on an NVIDIA Quadro RTX 8000 GPU (48GB) for around 40 epochs which took about five days.

An MSA subsampling strategy is applied during training, not only for regarding as data augmentation to train a robust model, but also for preventing the GPU from running out of memory when filled with large MSA. We randomly select MSA rows, up to a maximum of 2^14^/*c*, and down to a minimum of 16, though always including the query sequences in the first row. Large proteins of more than 960 residues long were discarded during training. We subsampled MSA with 256 sequences at the inference stage by adding the sequence with the lowest average hamming distance. We performed trRosetta protein structure modeling five times with the same input and selected the structure with the lowest energy for protein structure similarity measurement.

## Results

### Benchmark on long-range contact prediction

First, we benchmarked the long-range contact prediction performance of A-Prot using the FM and FM/TBM targets of CASP13 (Kinch et al., 2019). The benchmark results show that the performance of our model outperforms that of the existing methods. We compared the precision of our model’s top L/5, L/2, long-range contact predictions (long-range: sequence separation of the residue pair ≥ 24) and the other existing methods. The performance measures of the other methods are adopted from the reference (Wu et al., 2021). The top L/5, Top L/2, and L contact precisions of our model are 0.561, 0.712, and 0.813, which are higher than those of the other methods. For example, DeepDist, one of the state-of-the-art methods, predicted the top L/5, L/2, and L with precisions of 0.813, 0.712, and 0.517. Also, compared with AlphaFold predictions of CASP13, A-Prot predicted more accurately in all three measures by 7∼9%.

### Benchmark on model accuracy

In addition to contact prediction, we also compared the quality of protein models predicted by A-Prot with those submitted by the top-performing server groups of CASP14 (Table 2). First, we modeled the structures of 25 FM/TBM and TBM-hard targets of CASP14. The average TM-score and lDDT score of the models were compared with those of the following server groups: FEIG-S (Heo et al., 2021), BAKER-ROSETTASERVER (Anishchenko et al., 2021), Zhang-Server, and QUARK (Zheng et al., 2021). The model structures of the other groups were downloaded from the archive of the CASP14 website, and TM-score and lDDT scores were recalculated with the crystal structures and domain information for a fair comparison.

**Table 1.** Contact Precision on CASP13 FM and FM/TBM 43 domains (results adapted from DeepDist (Wu et al., 2021))

| Group | Top L | Top L/2 | Top L/5 |
| --- | --- | --- | --- |
| TripletRes | 0.451 | 0.587 | 0.700 |
| AlphaFold | 0.497 | 0.629 | 0.742 |
| RaptorX-Contact | 0.481 | 0.612 | 0.744 |
| trRosetta | 0.506 | 0.652 | 0.751 |
| DeepDist | 0.517 | 0.661 | 0.793 |
| A-Prot | **0.561** | **0.712** | **0.813** |

**Table 2.**
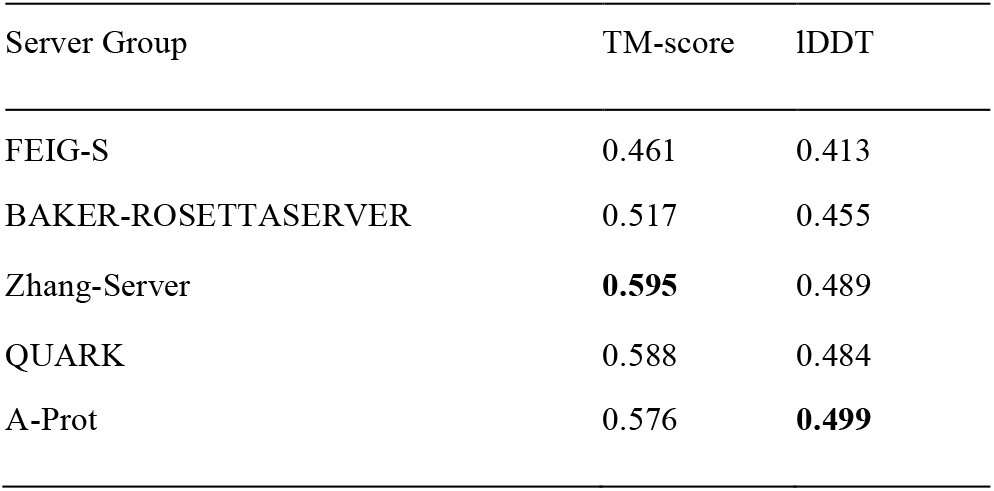
The average TM-score and lDDT of the model structures of 25 CASP14 FM, FM/TBM, and TBM-hard domains

A-Prot outperforms the other top server groups in terms of lDDT. In TM-score, A-Prot is showing slightly worse results, 0.576, than Zhang-Server and QUARK, whose TM-scores are 0.595 and 0.588. Compared with BAKER-ROSETTASERVER and FEIG-S, A-Prot models are consistently more accurate in both measures. These results show that the accuracy of A-Prot is comparable to or better than the top-performing server groups that participated in CASP14 (Kinch et al., 2021).

### A head-to-head comparison with ROSETTASERVER

Because A-Prot uses trRosetta for modeling structure at the final stage, we performed a head-to-head comparison of A-Prot models with the ROSETTASERVER models to identify improvement in residue-residue distance predictions (Figure 2). In terms of TM-score, many predictions made by A-Prot are significantly better than BAKER-ROSETTASERVER. For instance, the model qualities of five targets that were predicted to have TM-score less than 0.4 by ROSETTASERVER were improved higher than 0.4, corresponding to a correct fold prediction. Four highly accurate models are depicted in Figure 3.

**Figure 2.**
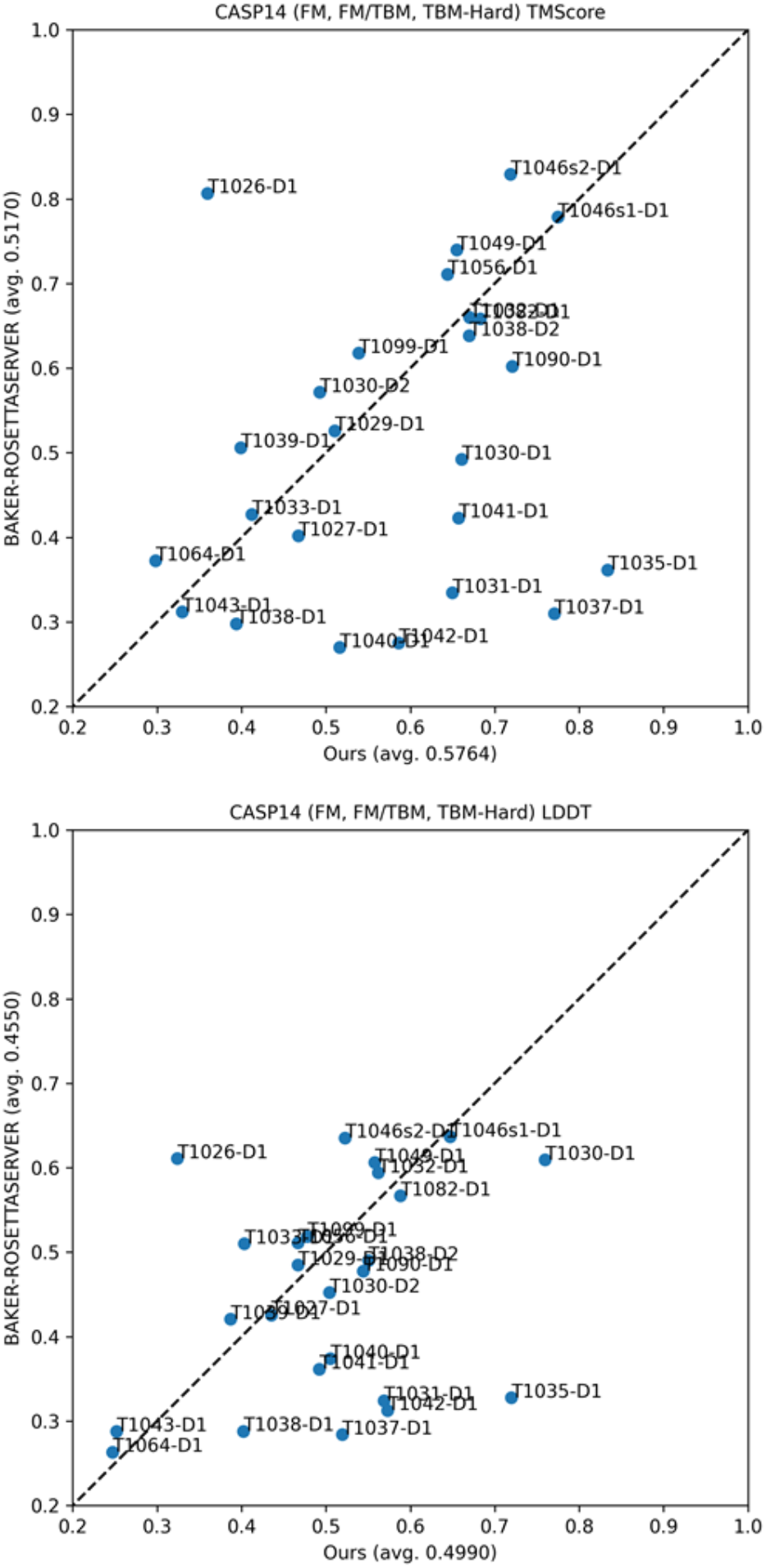
TM-score and lDDT on CASP14 FM, FM/TBM and TBM-hard 25 domains compared with BAKER-ROSETTASERVER.

**Figure 3.**
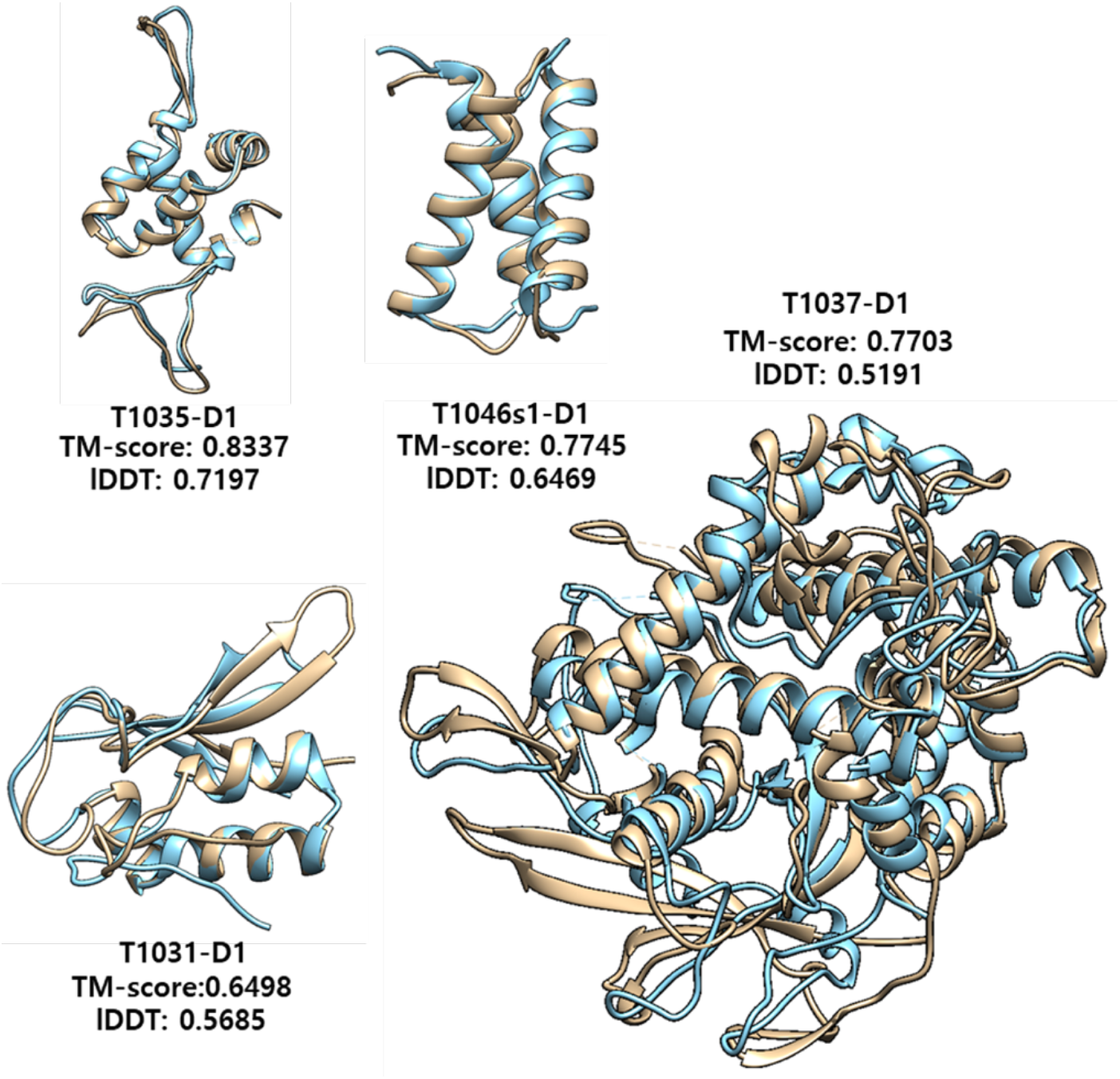
Model comparison of four high-quality CASP14 models generated from our method versus their native structures. Brown: native structure; Blue: model.

Similarly, in terms of the lDDT measure, prediction results of eight targets were improved significantly. On the contrary, only two targets deteriorated more than 0.05. Therefore, A-Prot predicts residue-residue distances and dihedral angles better than most server groups participated in CASP14 (Pereira et al., 2021).

On the other hand, nearly all worse predictions by A-Prot deteriorated only by a small margin except the T1026-D1 target. For T1026-D1, A-Prot did not predict its correct fold. This incorrect prediction is attributed to an incomplete MSA. T1026-D1 is a protein consisting of a virus capsid (PDB ID: 6S44). Our sequence search procedure found only less than 30 sequences, which appear to be not enough to extract correct evolutionary information. The failure of T1026-D1 suggests that having enough homologous sequences is critical in accurate 3D structure modeling using MSA-Transformer. In other words, a more extensive sequence search may improve the model accuracy of A-Pro. Except for T1026-D1, the deviations of all worse predictions than ROSETTASERVER were less than 0.1 TM-score, much smaller than the improvements.

## Conclusion

In this study, we introduced a new protein structure prediction method, A-Prot, using MSA Transformer. Our benchmark results on the CASP13 TBM/FM and FM targets show that A-Prot predicts long-range residue-residue contacts more accurately than the existing methods. We also assessed the quality of protein structure models based on the predicted residue-residue distance information. The model generated by A-Prot is more accurate than most of the server groups that participated in CASP14. The average lDDT of A-Prot models is higher than that of all server group models. In terms of TM-score, our model is slightly worse than QUARK and Zhang-Server. These results show that our approach yields highly accurate residue-residue distance predictions.

A-Prot requires less computational resources than the other state-of-the-art protein structure prediction models (Pereira et al., 2021). The source code of AlphaFold2 is only partially open (Jumper et al., 2021). Its model parameters are fixed, and only the structure modeling part is open. Thus, it is hard to tune AlphaFold2 for bespoken purposes. In addition, training the AlphaFold2 architecture requires a significant amount of computational resources, which is not assessable for most academic groups. On the other hand, A-Prot can be trained with a single GPU card. In summary, A-Prot will open new possibilities for training novel deep-learning-based models to predict various properties of proteins only using sequence information.

## Acknowledgment

This work was supported by the National Research Foundation of Korea (NRF) grant funded by the Korea government (MSIT) (No.2018R1C1B600543513).

